# A Tale of Two Hypotheses: Genetics and the Ethnogenesis of Ashkenazi Jewry

**DOI:** 10.1101/001354

**Authors:** M.A. Aram Yardumian

**Affiliations:** Department of Anthropology, University of Pennsylvania, Philadelphia, PA 19104-6398, USA; Department of History & Social Sciences, Bryn Athyn College, Bryn Athyn, PA 19009-0717, USA

**Keywords:** Caucasus, Khazar, haplotype, haplogroup, lineage

## Abstract

The debate over the ethnogenesis of Ashkenazi Jewry is longstanding, and has been hampered by a lack of Jewish historiographical work between the Biblical and the early Modern eras. Most historians, as well as geneticists, situate them as the descendants of Israelite tribes whose presence in Europe is owed to deportations during the Roman conquest of Palestine, as well as migration from Babylonia, and eventual settlement along the Rhine. By contrast, a few historians and other writers, most famously Arthur Koestler, have looked to migrations following the decline of the little-understood Medieval Jewish kingdom of Khazaria as the main source for Ashkenazi Jewry. A recent study of genetic variation in southeastern European populations (Elhaik 2012) also proposed a Khazarian origin for Ashkenazi Jews, eliciting considerable criticism from other scholars investigating Jewish ancestry who favor a Near Eastern origin of Ashkenazi populations. This paper re-examines the genetic data and analytical approaches used in these studies of Jewish ancestry, and situates them in the context of historical, linguistic, and archaeological evidence from the Caucasus, Europe and the Near East. Based on this reanalysis, it appears not only that the Khazar Hypothesis per se is without serious merit, but also the veracity of the ‘Rhineland Hypothesis’ may also be questionable.

---

There once was a book called *The Thirteenth Tribe*, written by a Hungarian Jew named Arthur Koestler. It wasn’t a very good book, inasmuch as its sources led short distances to nowhere, and considering plausible alternatives wasn’t on the author’s list of things to do. And yet its thesis—that Eastern European Jews are descended not from the Twelve Tribes of literary antiquity, but from Khazars, the shadowy Turkic stewards of a confederated multi-ethnic polity in the Medieval North Caucasus—has long outlasted the failure of the book’s dedicated purpose, which was to defeat anti-Semitism by claiming that European Jews weren’t Semites after all.^1^

That Judaism in some form was practiced by some element of the Khazar Empire in the 8^th^ to 10^th^ centuries CE is not in dispute (see Golb & Pritsak 1982 and Golden 2010, among many others). Whether and how this Jewish community became or influenced nascent Eastern European Jewry is the question. And, it is a question immediately bent by further uncertainties. Chief among them is how a group of Turkic- or Caucasian-speaking migrants could settle in Eastern Europe without introducing any discernible Turkic or Caucasian elements into Yiddish.^2^

In the absence of sufficient historical records, researchers have turned to population genetics as a platform for researching the broader brushstrokes of Ashkenazi Jewish ethnogenesis. Over the years, there have been numerous such studies, each with their own successes and limitations (Atzmon et al. 2010; Behar et al. 2003, 2004a, 2004b, 2006, 2008, 2010; Bray et al. 2010; Feder et al. 2007; Hammer et al. 2000; Kopelman et al. 2009; Listman et al. 2010; Nebel et al. 2001, 2005; Need et al. 2009; Shen et al. 2004; Skorecki et al. 1997; Zoossmann-Diskin 2010; Ostrer & Skorecki 2013). Last year, Dr. Eran Elhaik of the Johns Hopkins University School of Medicine made headlines with a study of European Jewish genetic structure that positioning the Khazar Hypothesis—which states that European Jewry finds its origins among Khazar converts and thus they share genetic affinities with neighboring populations in the North Caucasus and Southern Russian steppe—against the more traditional Rhineland Hypothesis—which asserts that European Jewry is descended from a small but expansive German Jewish population isolate, which in turn descended from Jews deported from Palestine to various parts of the Roman Empire between the first and seventh centuries CE. His conclusion was that the rise of Eastern European Jewry is explainable only by the Khazar Hypothesis.

Elhaik (2012) used a published data set of autosomal markers representing 1,287 unrelated individuals belonging to eight Jewish and 74 non-Jewish populations. Without being able to assume European Jews are the descendants of Israelite and Canaanite tribes, he had no obvious data set to use as a baseline. In addition, since there are no extant populations known to be directly descended from Khazar Turks or their imperial subjects, it was not self-evident as to whose genes should be used to represent this key reference group. Elhaik chose Palestinian Arabs as a proxy for Israelites, and modern day Georgian and Armenian populations as proxies for Khazar subjects. He made this selection because over 50% of Israeli Jews descend from the very population he seeks to contextualize, and the population of Palestine, European Jewish immigration aside, has not experience any statistically significant admixture in 2000 years (Haber et al. 2013; for a contrasting view, see Zalloua et al. 2008). Armenians and Georgians were chosen because of their proximity to the Khazar Empire, which occasionally included these regions within its boundaries.

Elhaik’s basic conclusion—that the Ashkenazi Jewish population is a mosaic of European and Middle Eastern ancestries—was hardly unexpected by linguists, historians, and others who research this subject. In fact, they corroborate notions that Eastern and Central European Jews share certain Middle Eastern and Western and Eastern European ancestries, the foremost of which was expected to be a contribution from Israelite migrants (or ‘exiles’), and the latter two from admixture with (or conversion from) non-Jewish Europeans over time. The distance of all Jewish communities from Middle Eastern populations—apparently the only thing shared by all Jewish communities—might also not be surprising. The controversy of this study, then, is its ostensible demonstration of similarities between European Jews and Caucasus populations, with a ‘specific Caucasus [maternal] founding lineage with a weak Southern European ancestry’ and a ‘dual Caucasus–Southern European [paternal] origin’ (Elhaik 2012: 69).^3^

The notion that Ashkenazi Jewry descends not from a Semitic population, however admixed, but from an entirely converted membership with no roots in Palestine would have strong and potentially hazardous ramifications for Zionist mythology. Not surprisingly, Elhaik has already sustained a barrage of attacks from the Zionist media. John Entine (who misspells Caucasus throughout his article) characterized Elhaik’s motivations as ‘more personal and ideological than scientific’ (Entine 2013). Seth J. Frantzman (who also misspells Caucasus throughout) says much the same (Frantzman 2013). Even Dore Gold chimed in, claiming the study was, or might be, ‘motivated mainly by a hostile political agenda that aims to advance the delegitimization of the Jewish state’ (Gold 2013). Many in the scientific community have also expressed distaste for the research. Marcus Feldman, Director of Stanford’s Morrison Institute for Population and Resource Studies, is quoted as saying, ‘If you take all of the careful genetic population analysis that has been done over the last 15 years… there’s no doubt about the common Middle Eastern origin’ (Rubin 2013).

And yet the question remains, how do we account for the sharing of specific ancestral markers among Eastern European Jews and Caucasian populations, ones which are otherwise unshared with other neighboring populations? Indeed, on the surface, it would be tempting to envision a westward migration of Judaized Khazars following the diminution of their empire (7^th^–11^th^ centuries CE) to explain this facet of Ashkenazi Jewish heritage.

However, there are several interrelated problems with the superimposition of Elhaik’s statistical results onto the Khazar Hypothesis that undermine this model for Ashkenazi origins. The first of these is that Armenians and Georgians are inappropriate proxies for the Khazar populace. Elhaik justifies their use with the assumption that ‘Khazarian, Armenian, and Georgian populations forged from [a confederation of Slavic, Scythian, Hunnic-Bulgar, Iranian, Alans, and Turkish tribes] were followed by relative isolation, differentiation, and genetic drift in situ’ (Elhaik 2012, p. 62).^4^

This statement is erroneous on three counts. To begin with, Armenians and Georgians are peoples indigenous to the South Caucasus, whereas the ‘Slavic, Scythian, Hunnic-Bulgar, Iranian, Alans, and Turkish’ influences to which Elhaik refers is almost exclusively limited to populations of the North Caucasus. While it is the case that North and South Caucasus populations do share very generalized ‘West Asian’ ancestry (see Figure 1), the distribution of genetic lineages, not to mention languages, suggests very different timings and patterns of settlement for the North and South (compare the results of Balanovsky et al. 2011 with those of Herrera et al. 2012 and Nasidze et al. 2004). In other words, populations such as Georgians, Armenians, Chechens, and Avars may share a Near Eastern ancestry in deep prehistory, but so do numerous other peoples of the Greater Near East and Mediterranean world, including possibly Palestinian Arabs, whose gene frequencies were used by Elhaik as proxies for ancient Judeans.

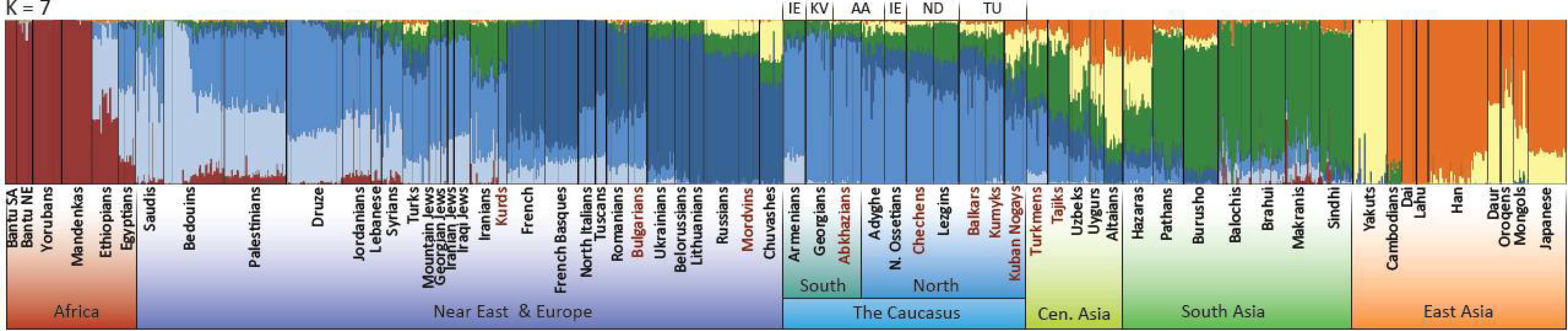

Furthermore, the nomadic Slavic, Scythian, Hunnic-Bulgar, Iranian, Alans, and Turkic tribes of the Southern Russian steppe (i.e., those most likely to have been subject to Khazar imperial subjection, whether they knew it or not) are demonstrably different in origin to their Caucasian neighbors to the south (Malaspina et al. 2003; Underhill et al. 2009; among other papers). Given this evidence, it is too much to assume that a South Caucasus population could stand as a proxy for Khazar imperial subjects in the North. Why, then, suggest these two populations were forged from the same ‘amalgamation of tribes’ as nomadic Turkic-speaking Khazars in the North Caucasus steppes?^5^

A second problem arises from predicating the statement about the forging of an amalgamation of tribes with the phrase ‘followed by relative isolation, differentiation, and genetic drift in situ’ (a paraphrase from Balanovsky et al. 2011, p. 2905). This assertion is inconsistent with geographic and chronological information for the region. In their paper, Balanovsky et al. (2011) are referring to population processes strictly in the North Caucasus, not the South. Secondly, it is a reference to concomitant processes that shaped the four major Y-chromosome haplogroups (G2a3b1-P303; G2a1a-P18; J1*-M267(xP58); and J2a4b*-M67(xM92)) and linguistic groups (Adygo-Abkhazian (i.e., Northwest Caucasian); Eastern Iranian; Nakh; and Daghestan), and thus began as far back as the Upper Paleolithic—millennia before Khazar imperial rule. The fall of the Khazar Empire and the subsequent age of Russian imperialism, in fact, signaled the beginning of the end of the ‘relative isolation, differentiation, and genetic drift in situ’, although even today the population bases for the North and South Caucasus remain divergent.

Moreover, it cannot be said that the South Caucasus was so integral to the Khazar Empire that Armenians and Georgians would be in any way biologically typical of it. Although both Armenia and Georgia at times became staging ground for wars fought by the Arabs against the Khazar armies, there is no mention in either Armenian or Georgian historiography of a period of Jewish conversion, or indeed of Khazar suzerainty, in either kingdom. Movsēs Xorenac’i’s *History of Armenia* and the *Kartlis Tskhovreba*, are the primary historical sources on this subject, and various later works (e.g., Lang 1966) cite them. Armenian and Georgian sources are scant on reports of Khazar activity, including the conversion to Judaism, and when they do appear, references to the fact of conversion, and to the Khazars themselves, are imbued with geographic distance.^6^

An additional problem inherent in the glossing of Elhaik’s statistical results onto the Khazar Hypothesis lies in the statement ‘Caucasus Georgians and Armenians were considered proto-Khazars because they are believed to have emerged from the same genetic cohort as the Khazars’ (Elhaik 2012, p. 64). Here again, Poliak (1951) is cited, this time along with Dvornik (1962) and Brook (2006). Aside from the foregoing reservations about source populations in the Caucasus, this statement further presumes knowledge of who the Khazars actually were. It is not entirely clear if Elhaik intends ‘proto-Khazars’ to refer to the Turkic-speaking tribes who conquered the North Caucasus steppe, or means the antecedents to the populace of the Khazar Empire in all its diversity. Surely an empire as large as Khazaria could not ever have been monoethnic, let alone biologically homogeneous. In either case, we know almost nothing about it, or who among the Khazar subjects converted to Judaism. Whether the original Turkic-speaking tribal elite, the trading class, citizens of Balanjar and Atil, or Kiev and the Crimea, we are no closer to finding an appropriate proxy since there are no written records concerning its demography.

Therefore, we are left not with a link between Eastern European Jews and Khazars, but between Eastern European Jews and Armenians and Georgians, which in itself warrants an explanation lest we are drawn into envisioning a South Caucasus-based origin for Eastern European Jewry. The explanation requires a deeper look at population genetics and ancient history and, though there are lacunae in each set of data, clarity is not out of reach. If Eastern European Jews share a substantial and unique portion of their genes with Georgian and Armenian populations—a portion otherwise restricted to these latter two groups—and to a lesser extent with Palestinian Arabs, but not with Eastern European populations at large—this may due in whole or in part to population processes beginning in the Levant, during the Neolithic, Bronze Age, Iron Age, and Byzantine eras.

It has been suggested many times (e.g., Ammerman & Cavalli-Sforza 1984; Battaglia et al. 2009) that the so-called Neolithic ‘wave of advance’ did not play an important role in the peopling of the South Caucasus, or even bypassed it altogether. Yet, settlement tells all over the South Caucasus, predominantly on the Georgian plains, appear to have been established between the 6^th^ and 4^th^ millennia BCE by what appear to have been extended families or oikoi who set up 10 to 15 km from one another (see Ессен 1963; Чубинишвили & Кушнарева 1967; ჯაფარიძე & ჯავახიშვილი 1971; Кушнарева 1974, 1977; Мунчаев 1975, 1982; Нариманов 1966, 1982). In Georgia, these settlements are associated with two distinct but geographically overlapping Neolithic cultures, the Shulaveri-Shomu (5500-4500 BCE) and the Sioni (c. 5^th^ – 4^th^ millennia BCE).^7^ Various ceramic vessels and implements from the earliest phases of the Shulaveri-Shomu bear strong resemblance to materials at Çatalhöyük (layer 5 and on), Hacilar, Hassuna, and Jarmo (See Кигурадзе 1986). For example, an antler sickle with a groove for holding inserts found in the lowest layers of Kyul-Tepe I strongly resembles another recovered at the Hacilar site (See Нариманов 1982, p. 25). The ceramic styles of the middle and late phase of Shulaveri-Shomu ceramics likewise resemble (and in some cases are identical to) those of the Neolithic cultures of the greater Near East, such as Çatalhöyük, Hacilar, Hassuna, and Ubaid (See Нариманов 1982, 1987, 1992). Affinities shared between Bronze Age pottery types in the Caucasus and eastern Anatolia, as well as between eastern Anatolia and the Levant (the Khirbet-Kerak ware of Jordan and Palestine, and Kura-Araxes wares of Georgia and Armenia being the most conspicuous example), are indicative of cultural, if not population, continuity between these regions.

From a genetic perspective, two studies have associated Y-chromosome haplogroups J1 and J2 with the spread of agriculture and domestication from the Fertile Crescent (Chiaroni et al. 2008, 2010), and a separate study (Balaresque et al. 2010) connects R1b to these events. These three haplogroups combined account for 65% of Armenian male ancestry (see Herrera et al. 2012), suggesting the region, while perhaps sparsely populated throughout the Paleolithic and Mesolithic, was settled by agriculturalists from the Levant, whose genes survive today. Moreover, the distribution of haplotypes within Y-chromosome haplogroup T, a further 8% of Armenian male ancestry, clusters entirely with Levantine individuals. Herrera et al. (2012) also position Armenians as descendants of Neolithic settlers in eastern Anatolia and the Ararat Valley.

Around the turn of the fourth millennium BCE—well before the advent of Maikop—distinct, non-local, high quality ceramic vessels begin to appear throughout western Georgia in a pattern along the Mtkvari (Kura) and Rioni rivers, leading to the possibility of a northern migration route from eastern Anatolia into western Georgia and the Maikop cultural zone (See Пхакадзе 1988; see also Pitskhelauri 2012). The sites along which we trace such a route begin with Ziyaret Tepe, Hanago, Aştepe, and Çolpan in eastern Turkey. Moving into Georgia, we find materials at the Qvirila Gorge, Samertskhle Cave, Samele Klde, Abastumani, Orchoshani, Dzudzuana Cave, White Cave, and Darkveti. While fine-grained population genetics research is lacking in both western Georgia and eastern Turkey, the abovementioned studies of gene flow indicate a Levant-to-Caucasus direction, not the reverse.

The ethnogenesis of Armenians is a subject not yet thoroughly understood. However, classical and epigraphic sources place a people with this name in southeastern Anatolia (in the general vicinity of Urartu, Assyria, Media, Persia, and Babylon) about 500 BCE. It has been pointed out that the Armenian endonym *hay* is the result of the loss of an intervocalic -t- (as would be characteristic) from the form *Hati-yos* (i.e., Hattian, a people on whose lands the Hittites came to settle in about 2000 BCE). Moreover, several Armenian terms imbued with religious significance share analogs in Hittite as well as Phrygian (see Russell 1997).^8^ Although the components of Armenian ethnohistory are undoubtedly variegated, the notion that they must have migrated en masse from the eastern Balkans has been, for the moment, relieved of its import since the publication of a studying making a claim for Anatolia as the homeland of proto-Indo-European (Bouckaert et al. 2012).

Although studies of Georgian Y-chromosome diversity are currently uneven and at low resolution, the predominant paternal lineages among Armenians (Hererra et al. 2012) are found in not unexplainably different proportions among Ashkenazi Jewish men (Behar et al. 2004; Hammer et al. 2009; Nebel et al. 2001; Semino et al. 2004; Shen et al. 2004):

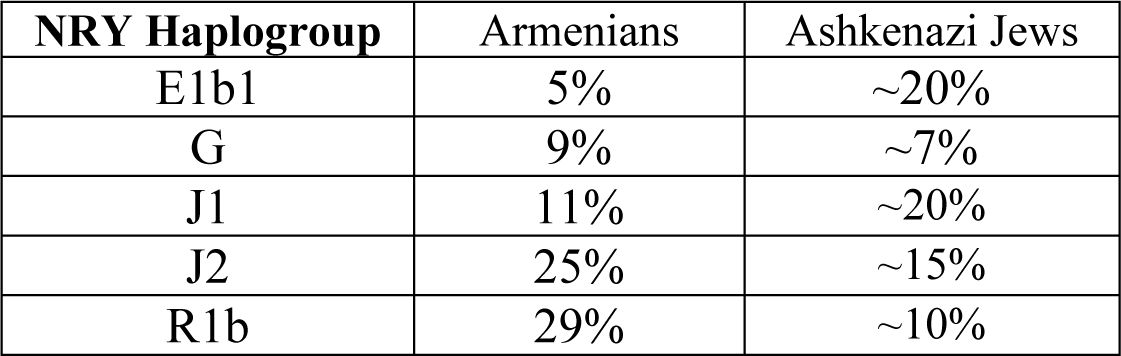

In fact, there are no major Y-chromosome haplogroups found among Ashkenazi Jews that are not also found among Armenians and vice versa, which would indicate the source populations for these groups, while diverse to begin with, were in geographic proximity.

Mitochondrial DNA (mtDNA) affinities vary somewhat more, perhaps due to the legality of matrilineal heritage among all Jews, as well as to the apparent European autochthony of these lineages (see Costa et al. 2013), but all the principal haplogroups are present (see Ottoni et al. 2011 for Armenian data; Feder et al. 2007: 499 for Ashkenazi data; tables for Romanian and Russian Ashkenazi Jews are also included there):

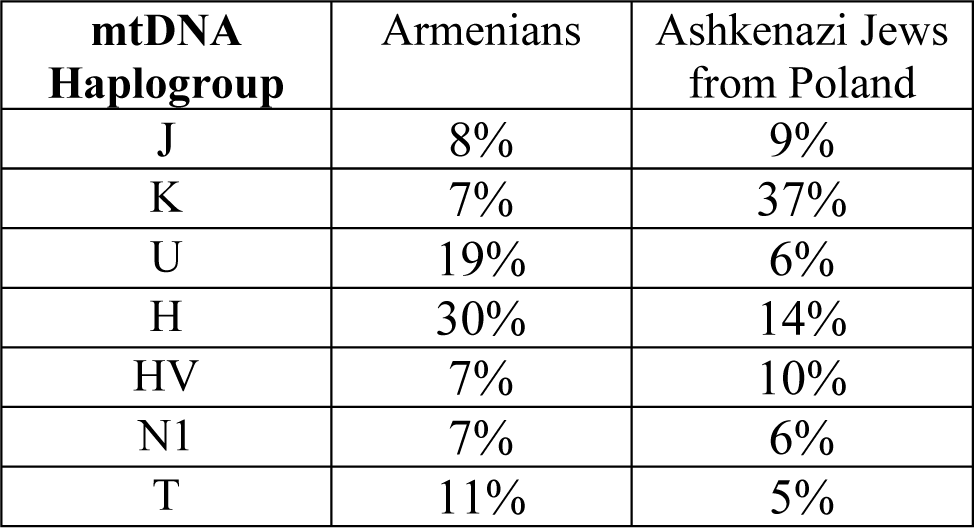

If a significant apportionment of genes are shared between Armenian and Jewish populations, and furthermore if this is due to the geographic proximity of their source populations, then, ironically, Elhaik may actually have helped confirm the narrative he was attempting to refute. Then again, the Khazar Hypothesis, as Elhaik comes to it, does leave room for the presence of Byzantine (specifically Anatolian) and Persian converts to Judaism in the gene pool (see Elhaik 2012, p. 62). Even if these populations came together in Eastern Europe rather than in Khazaria, this possibility alone could help resolve the question of what appear to be Caucasus-specific markers in Ashkenazi populations.

What, then, can we salvage from Elhaik’s study about Ashkenazi Jewish population histories? The biostatistical case demonstrating two distinct major components in European Jewish ancestry is very strong, regardless of the veracity of the proxies. The apparent absence of Caucasus-specific markers (as well as Middle Eastern-specific markers) in non-Jewish Eastern European populations (the Turkic-speaking Chuvash here serving as proxy) would seem to minimize the possibilities of Jews acquiring them before they arrived in Eastern Europe. Moreover, the apparent closeness of European Jews and Caucasus populations and the distance between these Middle Eastern populations would also seemingly indicate a Caucasus source for Eastern European Jews. But, since Middle Eastern and Caucasus ancestries are treated statistically as if separate, the proportional ancestry results are skewed. It is difficult to say exactly what the admixture analysis would look like had Middle Eastern, Near Eastern, and Caucasus populations been more accurately rendered.

Therefore, when historical migration trajectories are taken into account, the Khazar Hypothesis loses its viability, although not entirely so. Elhaik’s version of this hypothesis does allow for contributions from Greco-Roman Jewry as a way of explaining the patterns he observed in his data analysis. Since we have closed the gap between his ostensibly separate ‘Caucasus’ population and other populations of the Middle East, a migration of ‘Greco–Roman male-driven migration directly to Khazaria’ (Elhaik 2012, p. 69) is no longer a necessary part of the story. The observable patterns could just as well be explained by direct migrations of Jews into Central Europe from the circum-Mediterranean region and Russia. No population genetics study of Jews across Europe has ever indicated total homogeneity, and a singular Judean ancestry cannot account for the vast population of Eastern European Jews in the beginning of the 20^th^ century. Nor is anyone in doubt that Judaism continued to be a proselytizing religion well into the Middle Ages (See e.g., Bamberger 1939; Golb 1988; Goodman 1989; Rapaport 1965). The question, therefore, has always been what the apportionments of genes can tell us about patterns of Jewish settlement in Europe.

If a case for post-Khazar dispersal is to survive among other contributions to Jewish genetic diversity, then it will likely hinge on the presence of Y-chromosome haplogroup R1a1a, which has been reported as high as 20% among Ashkenazi men (Shen et al. 2004). This haplogroup is found quite infrequently among Middle Eastern populations, peaking instead in Western and Central Europe, and occurring at significant frequencies in South Asia. Its presence among certain Turkic-speaking populations^9^ and Kazan Tatars (Wells et al. 2001), as well as Finno-Ugric-and Slavic-speaking populations of the European Russian steppe,^10^ makes it a suitable focus for further analysis.^11^ However, if no trace of East Asian ancestry can be identified generally or in a specific cluster of Eastern European Jews, then the Khazar Hypothesis, as Elhaik presents it, is in even deeper trouble.

This brings us back to the apparent Greco-Roman Jewish contribution to the Ashkenazi gene pool. Whether or not they were Jewish proselytes, Khazars controlled trade along the Dnieper, Don and Volga Rivers—a portion of land wedged between Byzantium and the land of the Rus, the latter of whom eventually knelled death to the Khazar Empire around 965 CE. It is not, therefore, so difficult to imagine Hellenic and Roman Jews settling into trading posts all along the Dnieper and the roads leading into Baltic lands and Germany. In fact, the Solomon bar Simson Chronicle describes Crusader-era massacres (c. 1096) in the well-established Jewish communities of Mainz, Worms, Speyer, and elsewhere along the Rhine, as well as Trier, Regensburg, Mehr, and Prague, and in Bohemia (bar Simson 1977). Of potential sources for Jewish populations and the routes they may have taken into Eastern Europe and the Pale of Settlement, there are so many other possibilities worth considering.

The literature on Hellenistic Jewry is longer on textual criticism than on social details. Nevertheless, with the addition of archaeological sources, we can be sure Jewish communities in Asia Minor thrived with proselytes and converts. Whether these were voluntary or more compulsory, as was the case with the Edomites and Idumeans (see Josephus 15:9), it is unclear, but more likely the former given the lack of evidence for Jewish military apparatus in Asia Minor. Moreover, the well-known campaign to translate the Old Testament into the Greek dialect Koiné (which served as the lingua franca of much of the Mediterranean region following the imperial consolidation of Alexander the Great in the 4^th^ century BCE) must have been planned with converts in mind.

From Greek epigraphic and papyri sources, we know of scores of synagogues in Asia Minor,^12^ the northern Black Sea region,^13^ and southeastern Europe, chiefly in the Balkans, but also Hungary.^14^ Although we have no membership lists (or even statements of function) for these synagogues, a great many converted gentiles are mentioned in Rabbinical literature (Bamberger 1939: 221-266; also on this general subject see Shapiro 1960; Rapaport 1965). Baron (1952, p. 170 & 370-372) believes there were between four and eight million Jews outside Palestine in the first century CE. Even if the number was five million, this amounts to one-tenth of the population of the entire Roman Empire according to the censuses of Augustus (Durand 1974, p. 253-296). This ratio is unimaginable without the conversions of large numbers of gentiles. The question, then, is whether and to what extent the Jewish communities of Asia Minor, the Black Sea Region, the Balkans, and perhaps elsewhere, survived to influence Ashkenazim and how many succumbed to the pressures of rising state Christianity. Furthermore, it is not clear how many subsequently departed for central Europe in the waning days of Byzantium.

In the first centuries CE, the rate of conversion to Judaism in Rome was remarkably high (Bamberger 1968; Braude 1940; Rapaport 1965; Rosenbloom 1978). A diminished version of the Jewish Exile narrative would thus be that Ashkenazi Jews descended from these Roman converts as much as from the Judean immigrants or ‘deportees’ themselves. This is the opinion of Zoossmann-Diskin (2010), whose analysis of genetic markers on autosomes, sex chromosomes and the mtDNA positioned Eastern European Jews closer to their non-Jewish neighbors, especially Italians, than to other Jewish populations. Their genetic proximity to Italians led Zoossmann-Diskin (2010), as well as the authors of similar studies (Atzmon et al. 2010; Behar et al. 2010), to assert an origin for Ashkenazim among Roman converts to Judaism. Zoossmann-Diskin’s mistake was in thinking his modern day Italian proxies, and indeed ancient Romans, were not already the product of genetic interactions across the Mediterranean. In fact, when Neolithic settlement patterns in Southern Europe are taken into account (see Brotherton et al. 2013; Rootsi et al. 2004; Soares et al. 2010), they were certainly sharing genetic variation with their distant neighbors in Anatolia to the east (see Battaglia et al. 2009; Novembre et al. 2008).

Despite these issues, this paper, along with those of Atzmon et al. (2010) and Behar et al. (2010), represents a great improvement over the landmark study by Hammer et al. (2000) which calculates aggregated populations of Russians, Germans, and Austrians (a fictitious population in this sense) and moreover applies the χ^2^ test to the aggregates, which should be avoided in cases of multi-normal datasets (see Holt et al. 1980). In addition, populations from key corridors and centers of Jewish settlement, such as Belarus, Lithuania and Poland, were left out of the study, and so the results—favoring a Levantine origin for Eastern European Jews—may be skewed.

The conversion of the Kingdom of Adibene to Judaism in the first century CE might also be seen as an important transition in Jewish demography, if only because according to the letter to the envoy of ibn Shaprut, it was from or through Armenia, not via Black Sea Jews, that Judaism was brought to Khazaria (see Golb & Pritsak 1982, p. 130-31). The question then is whether to see this and other conversions to Judaism in western Eurasia as parcel to the game of alliance-building and politicking, for much the same reasons as Germanic, Slavic and Altaic tribes were turned toward Christianity, and others toward Islam in those centuries, or whether converts in Judaized regions were inculcated by immigrants from Palestine or other diasporic centers. For example, Josef ha-kohen claims the Arab invasion of Persia in 690 BCE was the impetus for a great many Persian Jews to migrate north into southern Russia (Hacohen 1971, p. 6-7). To wit, a mid-13^th^ – mid-14^th^ century Jewish cemetery was recently discovered near the village of Eghegis in Armenia’s Syunik Mars, although the origin and fate of this community remain unknown (Brown 2001).

There is some documentary evidence, obscure though it may be, for very early Jewish communities in various parts of Eastern Europe, particularly Pannonia and elsewhere in Roman Hungary. Besides the aforementioned evidence of synagogue architecture in Intercisa and Stobi, grave and monumental epigraphy, as well as religious objects, attest to their presence in the 2^nd^ or 3^rd^ century CE. For the most part, the evidence suggests that they were soldiers in the Roman Army. Patai (1996, p. 29) claims that their presence can be explained as a result of Sassanid Persian pressure against the eastern borders of the Roman Empire, causing Jews to flee to Pannonia, where they enjoyed full security, and presumably elsewhere in Europe. After the Romans abandoned Pannonia the fate of these Jewish lineages is unknown, for the first literary evidence for Jews in Hungary does not appear until 953 CE, with documentation of the arrival of a Croatian delegation in Cordoba.

The existence of Knaanic (a Western Slavic language related to Czech and Moravian, and thus of a Judeo-Slavic ethnos living east of the Elbe River) is indicative of a long presence in Eastern Europe, long enough at least for such a language to become its own, without apparently, a Hebrew substratum. Since several Medieval Hebrew texts (including some by Rashi) were glossed in Knaanic/Old Czech, this Jewish community must have been both observant and literate (Kearney 2010, p. 149, note 104). Linguist Paul Wexler has made a convincing case for Yiddish being a relexified form of Sorbian, and thus remains a Western Slavic language with High German vocabulary (Wexler 1991, p. 1–150, 215–225; 1993: 66–68). He subsequently extended this theory to include a parallel relexificative process in Kiev-Polessian languages (Wexler 2002). Knaanic (also called Judeo-Czech), which is a close relative of Old Czech and thus itself Slavic in origin, is another ethnolect which may have contributed to Yiddish. Moreover, around 1600, Rabbi Meir Katz Ashkenazi of Mogilev (in eastern Belarus) complained in a responsum on the subject of bilingualism that, ‘the majority of our coreligionists, who live in our midst, speak Russian’ (although by Russian it may be meant Russian per se or another Slavic language) (see van Straten 2011, p. 126, who, in turn, cites M. Katz Ashkenazi’s Hebrew/Aramaic-language book from 1687). Although little overall is known about Judeo-Slavic communities as they were, the fact that Yiddish, Slavic or Germanic contains so little of the Romance languages, so little Greek, and so little Aramaic or Hebrew, suggests that the word *ashkenaz* does refer to Scythia after all (see Wexler 2011, p. 277-78).

Jewish merchants and slave-traders abounded in antiquity all across Eurasia (see Hezser 2005). Perhaps the best documented of these are the Radhanites, a loosely confederated (if confederated at all) group of mobile multi-lingual Jewish traders who seem also to have been experts on route geography and social relations. Gil (1974) proposes, following Heyd (1885-86), that they were Arabic-speaking Jews from Radhan, a village that once existed in southern Iraq, and were successful not because of their incorporation but because of a network of Jewish settlements all across Eurasia, with whom they cooperated. Following the loss of their monopolies at the end of the 10^th^ century due to the rise of Genoa and Venice, and others whose sea-blockades and anti-Jewish campaigns destroyed their competition (Rabinowitz 1948, p. 188-90), they may have resettled in Mesopotamia, or moved on to Europe. However, what of the great network of Eurasian Jewish communities, particularly those in Russia, on whom they must have relied?

With so little Jewish historiography between Josephus Flavius (1st century CE) and the late 18^th^ century, it was natural for European historians to connect modern Jews directly to the ancient Judeans by way of a wholesale deportation to Rome. As Shlomo Sand points out, ‘Rome’s great Arch of Titus shows Roman soldiers carrying the plundered Temple candelabra—not, as taught in Israeli schools, Judean captives carrying it on their way to exile. Nowhere in the abundant Roman documentation is there any mention of a deportation from Judea’ (Sand 2009, p. 131). Nevertheless, this narrative has captured the imaginations of Jews and Christians alike because it depicts a small, resilient people bonded together in defiance of the Roman legions who would suppress their freedom to exist, thrive, and regenerate. The concept of a Jewish exile, as such, is critical to the concept of its return. Therefore, Zionist historians have in their scholarship minimized ideas relating to pre-Revolt diasporas, proselytism and conversion (in Rome and elsewhere), and voluntary departure from Palestine for reasons of mercantilism. And yet, this narrative of Exile has no more or less to recommend it than the various suppositions and rumors mentioned above.

Thus far, all major genetics studies, Elhaik’s excepted, have suggested that a majority of Jewish Y-chromosome genes are shared with other populations of the Middle East presumed to be autochthonous to the region. Nearly all these studies have concluded a relationship exists between the major diasporan Jewish, Ashkenazi, Sephardic, and Mizrahi populations. Y-chromosomal and mtDNA data, on the other hand, have indicated founder effects both of Middle Eastern and local origins, as well as significant admixture with local populations (Behar et al. 2003, 2004, 2006, 2008; Hammer et al. 2000; Nebel et al. 2001, 2005; Shen et al. 2009; Skorecki et al. 1997). Most significantly, there are apparently as few as four maternal haplogroups which make up ∼70% of Ashkenazi Jewish mtDNAs (K (32%), H (21%), N1b (10%), and J1 (7%). While it has been claimed these four lineages are of general Middle Eastern in origin (Behar et al. 2004, p. 4-5; see also Behar et al. 2006), the most recent research (Costa et al. 2013) suggests they are, by and large, mostly rooted in European prehistory, and their presence therefore must be due to conversion. Whole genome and autosomal studies (Atzmon et al. 2010; Bray et al. 2010; Kopelman et al. 2009; Need et al. 2009; Zoossmann-Diskin 2010) have bridged these seemingly divergent results by confirming the genetic proximity of Ashkenazi Jews to both Middle Eastern and European groups, concluding essentially that the Jewish genome is a series of geographical clusters tied loosely together. Overall, the closest genetic neighbors to most Jewish groups are Palestinian Arabs, Bedouins, and Druze (see Hammer et al. 2000; Nebel et al. 2000), southern European populations such as Cypriots and Italians (see Atzmon et al. 2010; Zoossmann-Diskin 2010; see also Behar et al. 2010), and, somewhat surprisingly, North Caucasus populations such as the Adygei (see Behar et al. 2003; Levy-Coffman 2005; Kopelman et al. 2009), the last finding being what inspired Elhaik’s endeavors to revisit the Khazar Hypothesis.

As to the issue of whether Jewish peoples are coreligionists—the aggregation of Judeans with Greco-Roman and possibly Slavic converts, or a single people, the descendants of a tribe with its roots in Palestine (see Entine 2007 and his other publications)—this remains unresolved because too much of the Ashkenazi Jewish genome is shared by other eastern Mediterranean populations. The current evidence does indicate ‘Middle Eastern’ origins, however admixed, for Ashkenazi Jewish male lineages, but how much does this really mean? To conclude that any population is Middle Eastern by descent is really to say very little, even if the proportional ancestry clusters join geographic neighbors (Figure 1). The Middle East is a big place and seems to have served as an incubator for human genes between the anatomically modern human departures from Africa and various later departures into Eurasia during the Upper Paleolithic, Mesolithic and Neolithic eras. Many of its currently visible phylogeographic gradations appear to be due to recent cultural barriers to gene flow (Haber et al. 2013), while the majority of Eurasian mtDNA and Y-chromosome haplogroups not directly associated with East Asia and India have their origins there, very generally. Given both these facts and uncertainties, why do we need either Khazaria or the Rhineland, specifically, to serve as geographic interfaces for Ashkenazi ethnogenesis, when, just as likely, the process was complex, gradual, and inclusive of converted Jews from a wide variety of lineages?

If *The Thirteenth Tribe* is better placed in the fiction section, maybe the Rhineland Hypothesis also belongs there. However, there is a further, more pressing question to be asked: why does the proposal of alternate hypotheses for the ethnogenesis of Jews create such alarm? Why does Dore Gold write of Elhaik’s research as aimed at advancing ‘the delegitimization of the Jewish state’? After all, no genetics study has ever isolated Palestine as a point of origin for Jewish populations, and the legitimacy of the state of Israel has nothing to do with genetics per se. Given this, the Zionist critics need not worry, unless, of course, there is population genetics research aimed at legitimizing Israel or any other revanchist project. Such studies would indeed be cause for alarm. Indeed, perhaps the wording of the recognition of a Jewish ‘race’ at Paris Peace Conference in 1919 and its relationship to the mandate for a creation of a Jewish state in Palestine should be studied more carefully.

There are numerous notions about the origins of all peoples, some parsimonious, some controversial, some purely mythological. Although much high quality genetics research points to a commonality of certain lineages within the Ashkenazi Jewish population structure, far too much is assumed in stressing Palestine as the point of geographic origin. To make finely pointed suggestions about the geographic origins of haplotypes amounts to guesswork, especially in the Middle East. Given the genetic structures of Greeks or Italians, one could construct a similar hypothetical argument for their origins in the region, and yet we know from history that there is more to the story. Without ancient DNA linking ancient Judeans to modern Jews, we can only position Ashkenazi Jewry beside their closest neighbors, which include the peoples of Anatolia, who for a time, it appears, saw more Judaized Hellenes than Hellenified Jews (See Levine 1998; Rapaport 1965; Rosenbloom 1978). Given the centuries of proselytic activities all around the Roman Empire, and perhaps the same in Persia and in southern Russia, it is a shame to see the story of Ashkenazi Jewish ethnogenesis cast in the terms of reductive narrative traditions such as Rhineland and Khazaria, rather than in a more complex and expansive framework.

## Acknowledgments

The author wishes to express thanks to Prof. Theodore G. Schurr, Prof. Aaron Brody, Dr. Miguel Vilar, and Akshay Walia for their suggestions for improvement to this paper.

## Footnotes

1 Koestler wasn’t the originator of the Khazar Hypothesis. He simply popularized ideas which had always in essence been in currency, but which had misfortunately been circulated by tendentious 19^th^ century racialists such as Ernest Renan and Lothrop Stoddard. Koestler himself seems to have drawn his conclusions primarily from his readings of AN Poliak (1941, 1951) and Hugo Kutschera (1910).

2 Paul Wexler (1990, 1993, 2002) has argued convincingly that periphrastic conjugation, a Yiddish linguistic feature held in common with Turkic languages and with Eastern Slavic languages, but not with German or German-derived slang lexicons that utilize Hebrew loan features via Yiddish, evidences Turkic influence on Yiddish at an early stage. His ideas about Ashkenazi ethnogenesis are among the most nuanced and useful.

3 Interestingly, if the data support a Khazar root in Eastern European Jewish ethnogenesis, they do not support a westward migration from the Caucasus to both central and southern Europe, as this would have been visible as a gradient from the Caucasus toward Europe for both matrilineal and patrilineal lines. Instead, Elhaik prefers to explain the observed pattern as a movement of Judaized Greco-Roman males from southern Europe to the Caucasus in the 6^th^ - 13^th^ centuries, followed by a movement west into eastern and central Europe in the 13^th^ - 15^th^ centuries.

4 For the amalgamation of tribes, Elhaik cites Poliak 1951, a book also heavily cited by Koestler. Since it is written in Hebrew I cannot comment on it or whatever sources it may cite in turn, but the statement is facile at best. For isolation, differentiation and drift, he cites Balanovsky et al. (2011), a high quality study.

5 Both the *Kartlis Tskhovreba* and the Armenian historian Movsēs Xorenac’i do, however, record the arrival of Jews in the region (which may also have included eastern Anatolia) after the fall of Jerusalem in 587 BCE. According to the Schechter Letter, Jews from Armenia and Persia comingled with nomadic Khazars, eventually assimilating (see Golden 2007 for a summary). One of the descendants of these settlers, named Sabriel, became a Khazar chief and was convinced by his wife Serakh to convert to Judaism. (Other sources credit Yitzhak ha-Sangari with having converted the Khazars to Judaism). These facts, alone or together, are worth consideration, but the use of today’s South Caucasian populations as surrogates for Khazar imperial subjects is highly problematic. More may be said about Khazar influence on the peoples of the North Caucasus, especially the Alans, about whom at least two sources recollect some amount of conversion to Judaism. But, here again, even if we consider North Caucasus populations as a more reasonable stock proxy for those Khazar subjects who converted to Judaism, the Caucasus Mountains have permitted very limited gene flow from the south (see Yunusbaev et al. 2011) and it was populations from the south which Elhaik has used.

6 See Thomson 1996, esp. pp. 5-6 wherein Khazaria is mentioned as a land in the north, beyond the mountains; pp. 13-16 wherein is described the Khazar invasion of the South Caucasus from the north is and its subsequent reversal by a Persian invasion from the south; p. 20 whereon refugees from Khazaria are mentioned as welcome in Georgia for their help against the Persians; p. 70 whereon Khazars from north of the Caucasus are called upon again for assistance in a war against the Persians; p. 78 whereon it is described the Khazar siege of Daruband [Derbent] from the north en route to invade Persia; p. 196 whereon Khazaria is clearly stated as a land distinct from Georgia; pp. 257-8 wherein the extension of Khazar sovereignty into Abkhazia the ruination of Tbilisi by Khazars are mentioned; p. 261 whereon it is described the settlement of 300 Khazar households in Shamkor (present-day Azerbaijan).

7 As yet, there are no C14 dates for Sioni sites.

8 During the ethnic cleansing of Canaan by the Israelites, as depicted in Joshua 3 (and presaged in Exodus 34:11 & 23:23, and Deuteronomy 7:1-3), one of the seven populations to be removed was the Hittites (the others being the Girgashites, Amorites, Canaanites, Perizzites, Hivites and Jebusites). While it remains unclear to what kind of social category the other terms in this list refer (whether linguistic group, urban/rural dichotomy, occupational caste, or something else), the biblical term ‘Hittite’ is by no means the same Hittites as the Bronze Age people of Anatolia who spoke an Indo-European language, and sold chariots to their Semitic neighbors to the south (see Singer 2006).

9 R1a1a is present in 27% of Karachays, 25% Balkars, 13% among Kumyks, 41% among Altaians (Underhill et al. 2009).

10 This haplogroup occurs at frequencies of 56% in Poles, 50% in Ukrainians (see Underhill et al. 2009); 37% in Udmurts (Semino et al. 2000), 51% in Belorusssians (Behar et al. 2003), 56% in Hungarians (Battaglia et al. 2008), 58% in Tver Russians, 52% in Kursk Russians, and 15% in Khanti (Mirabal et al. 2009).

11 A large-scale genomic survey of Eurasia west of the Urals and north of the Caucasus, perhaps using the POPRES dataset (e.g., Nelson et al. 2008; Novembre et al. 2008), might allow for a more complete demographic picture of these populations, the timings of their admixtures, and the dates for the most recent common ancestors (MRCAs) of Eastern European Jewry’s source populations among them.

12 Acmonia, Amastris, Ephesus, Hyllarima, Miletus, Myndos, Nysa, Pergamum, Philadelphia, Phocaea, Priene, Sardis, Side, Smyrna, Teos & Tralles, among others (see Kraabel 1979 & Feldman 1996).

13 Gorgippia, Olbia, Panticapaeum, Phanagoria, Sebastopol, among others. Op cit. note 9.

14 Intercisa & Stobi; Mursa & Oescus. Op cit. note 9.

